# Bacterial cell lysis: geometry, elasticity, and implications

**DOI:** 10.1101/343350

**Authors:** Felix Wong, Ariel Amir

## Abstract

Membrane lysis, or rupture, is a cell death pathway in bacteria frequently caused by cell wall-targeting antibiotics. Although several studies have clarified biochemical mechanisms of antibiotic action, a physical understanding of the processes leading to lysis remains lacking. Here, we analyze the dynamics of membrane bulging and lysis in *Escherichia coli*, where, strikingly, the formation of an initial bulge (“bulging”) after cell wall digestion occurs on a characteristic timescale as fast as 100 ms and the growth of the bulge (“swelling”) occurs on a slower characteristic timescale of 10-100 s. We show that bulging can be energetically favorable due to the relaxation of the entropic and stretching energies of the inner membrane, cell wall, and outer membrane and that experimentally observed bulge shapes are consistent with model predictions. We then show that swelling can involve both the continued flow of water into the cytoplasm and the enlargement of wall defects, after which cell lysis is consistent with both the inner and outer membranes exceeding characteristic estimates of the yield areal strains of biological membranes. Our results contrast biological membrane physics and the physics of thin shells, reveal principles of how all bacteria likely function in their native states, and may have implications for cellular morphogenesis and antibiotic discovery across different species of bacteria.

---

Antibiotic resistance is one of the largest threats to global health, food security, and development today.^1^ Its increasing prevalence^2^ begs the question of whether physical principles, which may be more universal than particular chemical pathways, could inform work on novel therapeutics, as has been done for mechan-otransduction in eukaryotes^3^ and tissue growth and fluidity.^4,5^ To elucidate such principles, a physical understanding of the cell death pathway caused by many antibiotics, which may complement knowledge of related biochemical mechanisms,^6,7,8,9,10,11,12^ is needed.

In many bacteria, cell shape is conferred by the cell wall, which resists the internal turgor pressure and is composed of two or three-dimensional layers of peptidoglycan (PG).^13,14,15^ In Gram-negative bacteria such as *E. coli*, the two-dimensional cell wall is sandwiched between the inner and outer membranes (IM and OM), while in Gram-positive species the cellular envelope comprises an inner membrane enclosed by a three-dimensional cell wall. PG consists of rigid glycan strands cross-linked by peptide bonds and is maintained through the combined, synchronized activity of enzymes including transglycosylases and transpeptidases.^13,15,16,17^ Many antibiotics, including penicillin and *β*-lactams, bind to transpeptidases to inhibit cross-linking. Inhibition of peptide bond formation, combined with mislocalized wall degradation by PG hydrolases, has been thought to result in large defects in the cell wall which precede bulging of the IM and OM and eventual cell lysis.^6,7,18,19^

## Results

### Dynamics of bacterial cell lysis

Inspired by previous work,^16^ we degraded wild-type *E. coli* cell walls with cephalexin, a *β*-lactam antibiotic, and observed typical cells to undergo the morphological transitions shown in Fig. 1A-C and Supplementary Video 1. Bulging—defined here as the development of a protrusion under an approximately constant volume, which is accompanied by a noticeable shrinking of the cell length—was observed to occur on a timescale as fast as 100 ms. Swelling, defined here as the growth of the protrusion, which may involve more variation in cytoplasmic volume, was observed to occur on a timescale of 10-100 s (Fig. 1D).^16^

**Figure 1:**
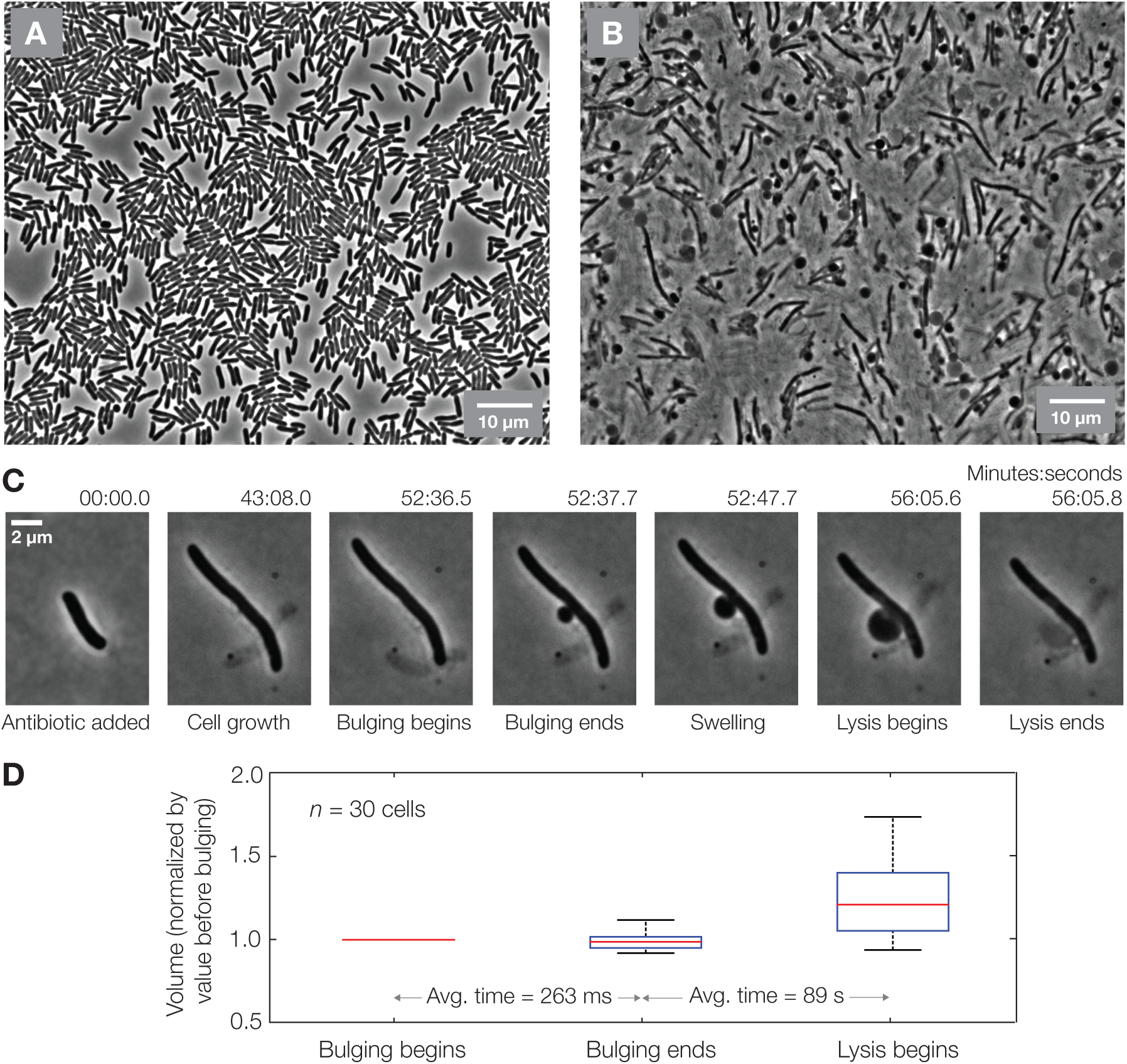
Experimental observation of membrane bulging, swelling, and lysis. (A) Phase-contrast image of a population of *E. coli* cells immediately after antibiotic treatment (see also Supplementary Video 1). (B) Phase-contrast image of the same population approximately 1 hr after antibiotic treatment, showing that membrane bulging and swelling is common to all cells (see also Supplementary Video 1). (C) Phase-contrast timelapse of a single *E. coli* cell during antibiotic killing, with the corresponding morphological dynamics denoted. (D) Image analysis of 30 bulging and swelling cells of varying lengths reveals that bulging usually does not result in a significant change in cell volume, while swelling is usually accompanied by more variation in volume. Bulged cells were fit to cylinders with protruding ellipsoids; see *Materials and Methods* for details on the image analysis methodology. Error bars represent variability between cells.

Physical modeling can elucidate the mechanisms which drive membrane bulging and swelling, and, as suggested above, understanding this is important because mechanical failure of the cell wall is a poorly understood, yet typical and exploitable, cell death pathway.^16,18,19,20,21^ Furthermore, the role of the cellular envelope here may entail different physics than that of eukaryotic blebbing.^22,23,24,25,26,27^ In a recent modeling study,^28^ a critical pore size for bulging was found by studying the trade-off between the bending energy cost of bulging and the pressure-volume energy gained. This trade-off appears to be irrelevant for determining bulge size in the aforementioned *β*-lactam experiments, where it can be shown that the bending energies are negligible compared to the stretching energies and that shortening of the cell contributes to bulge growth (Fig. 1C-D). As we shall see, membrane remodeling and the relaxation of the entropic and stretching energies of the cell envelope can predict bulging and are consistent with experimental observations.

### Cell envelope mechanics

We model the cell wall, IM, and OM as elastic shells in contact. Although fluid membranes cannot support in-plane shears,^29^ we do not consider shear strains and stresses in this work and the strain energy is effectively that of an elastic shell. Importantly, we also suppose that, on timescales longer than that of the elastic response, the membrane geometries can vary due to membrane fluidity while conserving their reference surface areas. This contrasts with the rigid cell wall, whose reference configuration is assumed to be a cylinder. The free energy of the cell wall, IM, OM, and the volume enclosed by the IM is

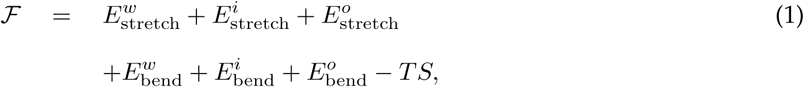

where the superscripts ^*w*^, ^*i*^, and ^*o*^ denote wall, IM, and OM quantities, respectively, E_stretch_ and E_bend_ are the stretching and bending energies, respectively, of an elastic shell, *T* is the temperature, and *S* is the entropy of mixing water and solutes. Here only water molecules are assumed to be outside the cell and *S* = −*k*(*n_s_* ln *x_s_* + *n_w_* ln *x_w_*), where *k* is Boltzmann’s constant, *x_s_* and *x_w_* are the number fractions of solute and water molecules, respectively, and *n_s_* and *n_w_* are the numbers of solute and water molecules, respectively. We assume an ideal, dilute solution in this work and note that the presence of the entropic term implies that, when the chemical potentials of water are equal both inside and outside the cell, the mechanical stresses in the cellular envelope are proportional to *p* = *kTC*, where *C* is the number density of solutes inside the cell and p is defined as the turgor pressure (Supplementary Information, SI). Assuming characteristic parameter values, the bending energies are negligible compared to the stretching energies, as is typically the case for thin shells.^30,31,32^ We therefore discard the bending energies in the expressions below and verify in the SI that they do not change our results. We assume linear, isotropic constitutive relations for the IM and OM and an orthotropic constitutive relation for the cell wall, building on evidence for a larger elastic modulus in the circumferential direction than the axial direction.^33,34^ *E*_stretch_ can then be expressed as 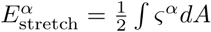, where *dA* is an area element and

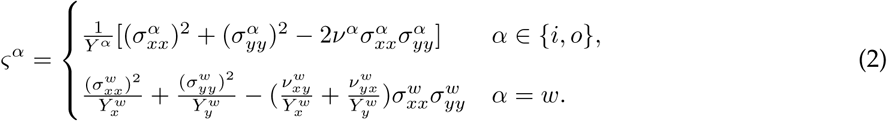

Here (*Y*, *ν*) are the two-dimensional Young’s modulus and Poisson’s ratio of the membranes, 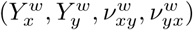 are the orthotropic analogues for the cell wall, and 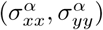 denote in-plane stresses in the axial and circumferential directions, respectively, of the *α* component of the cellular envelope. We relate *Y^i^* and *Y^o^* to the area-stretch moduli *K_a_* of lipid bilayer membranes by 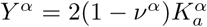. Values of *K_a_* have been estimated to be in the range of *K_a_* ≈ 0.03-0.25 N/m for *E. coli* spheroplasts depending on external osmolarity and size^35^ and *K_a_* ≈ 0.2-0.4 N/m for red blood cells (RBCs) and giant unilamellar vesicles,^36,37^ and these values are expected to be similar for bacterial membranes.^35^

As the membrane stresses may vary due to the in-plane rearrangement of phospholipids,^38,33^ we assume that they can be determined in a healthy cell by minimizing 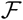 over the possible reference states. The existence of excess membrane area does not significantly change our results and involves calculations similar to that considered below. For a range of parameter values believed to be relevant to the *E. coli* cellular envelope, 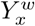 and 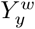 are similar in magnitude to *Y^i^* and *Y^o^*. Because 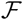 is quadratic in the in-plane stresses, the minimization of 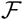 in Eq. (1) predicts that both the cell membranes and the cell wall are load-bearing. In particular, over the cylindrical bulk of a spherocylindrical cell for which 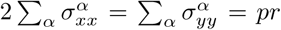 with *α* ∊ {*i, o, w*} and for characteristic values of 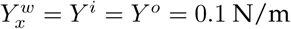,^35,33,39,40^ 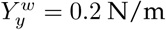,^33,34^ and Poisson’s ratios all set to 0.2,^41,42^ we find that 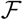 is minimal when 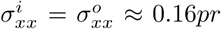, 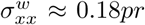, 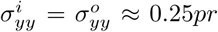, and 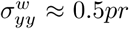, where *r* is the cell radius. This result contrasts with the idea that the cell wall is the only load-bearing structure of the cellular envelope and is consistent with experimental observations suggesting that the IM and OM can also be load-bearing, as manifested by the known fact that bulging precedes lysis.^16^ As the IM and OM are fluid, load-bearing by the IM and OM does not contradict the fact that *E. coli* cells become spherical without their cell walls.

When 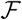 is minimized over the cellular dimensions, flow of water into the cytoplasm may be required. The bulk flow of water from the external milieu to the cytoplasm is thought to be characterized by the hydraulic conductivity *L_p_*,^43^ defined so that the instantaneous volumetric flow rate through a membrane is *dV*/*dt* = *L_p_A*_tot_*p*, where *p* is the turgor pressure and *A*_tot_ is the total membrane surface area.^43,44^ Estimates of *L_p_* vary depending on membrane structure: studies involving osmotically shocked bacteria,^45^ liposomes with aquaporin-1, and RBCs have found *L_p_* ≈ 10^−12^ m^3^/N · s, while studies for liposomes and other bilayers without water channels have indicated *L_p_* ≈ 10^−13^ m^3^/N · s.^43,46^ Below, we find that the larger value of *L_p_* is consistent with a volume increase on the order of 1-10% of the initial cell volume during bulging and that water flow can also contribute to swelling.

### Model of bulging

We now show that, over a timescale of 100 ms, removal of a piece of cell wall can result in bulging. As the amount of excess membrane surface area is finite,^35^ we assume that the surface areas of the reference states of the IM and OM remain unchanged over the timescale of bulging. We therefore consider a quasi-equilibrium state in which the membrane reference surface areas limit bulging, for which the envelope stresses correspond to those caused by turgor pressure loading. Furthermore, since osmoregulation is believed to occur on a timescale of ~1 min for osmotic shocks applied over less than 1 s,^44,47,48^ we assume the number of solute molecules to remain constant after cell wall degradation. The free energy may be lowered by water flow and bulging if the IM and OM may assume arbitrary geometries. Hence, we wish to minimize 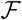 over the cell geometry and the cellular dimensions, assuming that the membrane reference surface areas are fixed.

Suppose that an area *A* of the cell wall is removed. For simplicity, we assume *A* to be a circle of radius *r_d_*. If the IM and OM were at equilibrium without bulging, then 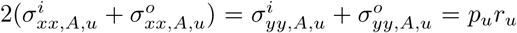; we denote the stresses satisfying force balance over *A* with the subscript *_A_* and quantities of the unbulged state with the subscript *_u_*. As discussed below, the solutes may be diluted due to water flow, so that *p_u_* = *pV*_cell_/*V_u_*, where *V*_cell_ = 2*πrL* is the volume of a healthy cell, *L* is its length, and *V_u_* is the volume of the unbulged state. Assuming the linear strain-displacement relations 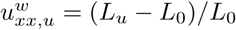 and 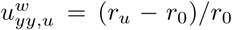 for the cell wall, where (*u_xx_*, *u_yy_*) denote in-plane strains and (*L*_0_, *r*_0_) the reference length and radius, respectively, the dimensions (*L_u_*, *r_u_*) of the cylindrical bulk in the unbulged state, which may differ from the dimensions (*L*, *r*) of a healthy cell, completely determine the stresses in the cell wall. The condition of force balance then constrains the stresses in the IM and OM over the remainder of the cellular envelope. Thus, the free energy of the unbulged state is

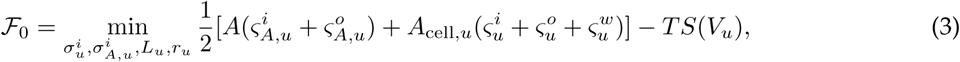
where 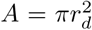 is the area removed, *A*_cell,*u*_ = 2*πr_u_L_u_* − *A* is the remaining surface area, ignoring the end-caps, *S*(*V_u_*) is the entropy of mixing corresponding to 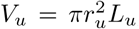, 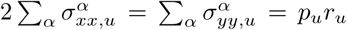 with *α* ∊ {*i, o, w*}, and the in-plane stresses are compactly denoted by *σ* = (*σ_xx_*, *σ_yy_*). (*L_u_*, *r_u_*) are related to (*L*, *r*) by the condition of fixed reference membrane areas, which for simplicity is considered only for the IM in this work and requires 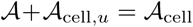, where the scripted symbols denote corresponding reference quantities and 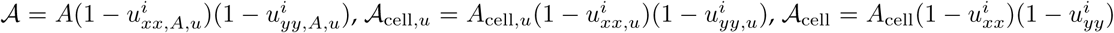, and *A*_cell_ = 2*πrL* is the surface area of a healthy cell. Due to water flow, the volume enclosed by the IM may increase, and we show below that the amount of volume increase is consistent with the timescale of bulging. We now wish to find cell envelope geometries for which the free energy is lowered.

As an ansatz which will later be supported by comparison with experiments, we suppose the formation of an ellipsoidal bulge with radii (*a, a, b*) and some circular cross-section coinciding with *A*, as described by the parametric angle *θ* = sin^−1^(*r_d_*/*a*) (Fig. 2A). We neglect the bending energy of the neck, as detailed in the SI (see also Fig. S1). As the reference membrane surface areas are conserved, bulge formation requires area to be appropriated from the cylindrical bulk. Below, we examine the change in stretching energy of the cylindrical bulk under the assumption that a reference membrane area 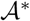 is removed and relegated to the bulge, and then include the change in stretching energy due to the formation of an ellipsoidal bulge over *A* with reference area 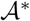 (whose calculation for an ellipsoidal shell is discussed in the SI).

**Figure 2:**
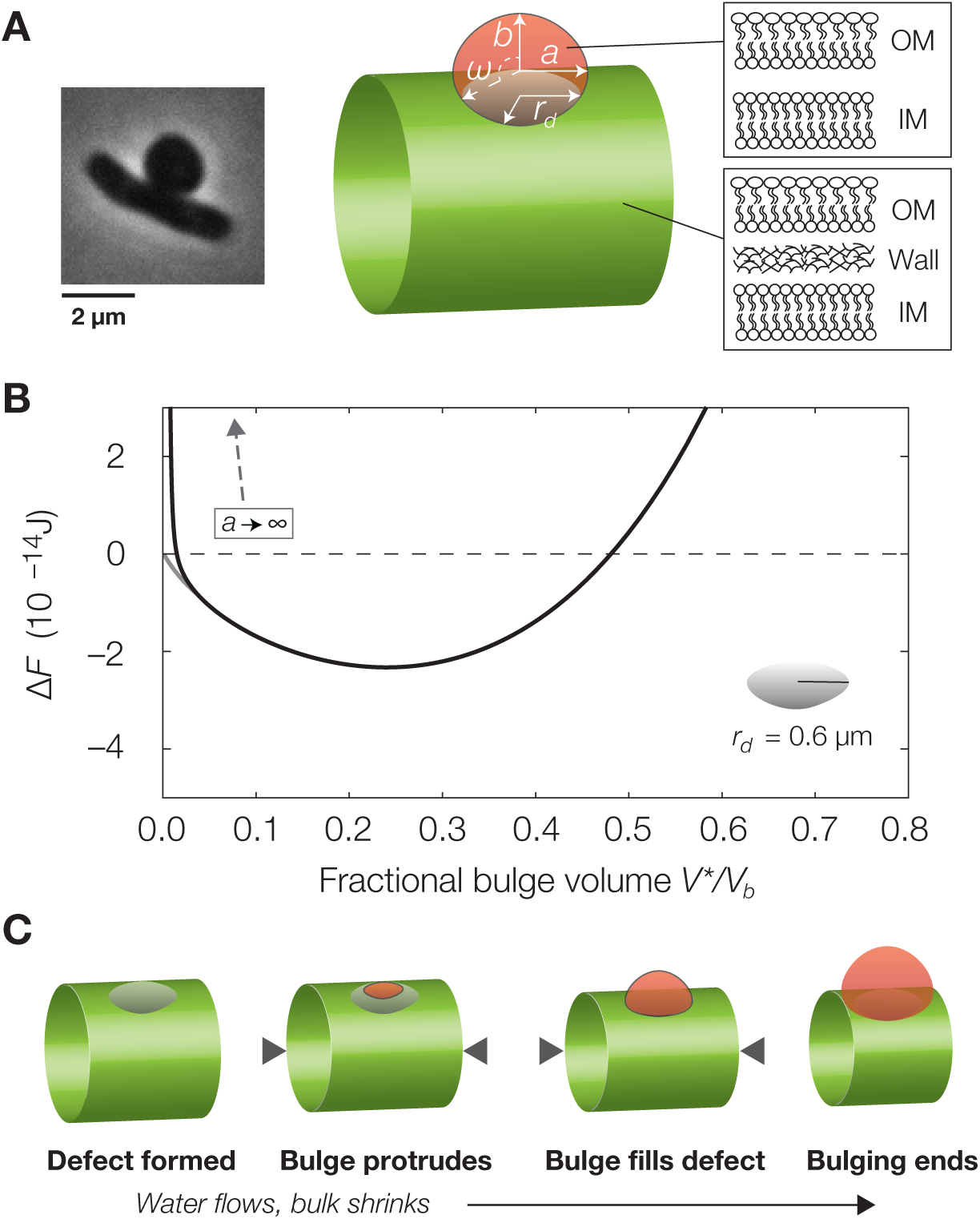
Energetics of bulging. (A) Schematic of a bulging cell. The subtended angle, *ω*, is related to *θ* by cos *ω* = *a* cos *θ*(*a*^2^ cos^2^ *θ* + *b*^2^ sin^2^ *θ*)^−1^/^2^. (B) Plot of 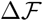 against the fractional bulge volume *V*^*^/*V_b_*, for the parameter values summarized in *Materials and Methods*. Here *V*^*^ denotes the bulge volume and *V_b_* denotes the cell volume in the bulged state. As an artifact of the ansatz geometry, 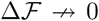 as *θ*, *V*^*^ → 0 because the bulge stresses diverge as *α* → ∞. Small bulges with diverging radii of curvature correspond to energetically unfavorable modes, but do not necessarily present energetic barriers, and the grey curve illustrates an alternate path to the energetic minimum which is monotonically decreasing in the free energy (SI and Fig. S2). (C) A possible relaxation path, for *r_d_* = 0.6 *µ*m.

To determine the change in stretching energy of the cylindrical bulk once a reference area 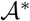 is removed, we first examine how the bulk contracts. As above, the dimensions (*L_b_*, *r_b_*) of the cylindrical bulk after bulging completely determine the stresses in the cell wall, and the condition of force balance constrains the stresses in the IM and OM. The stretching energy of the cylindrical bulk after bulging can be expressed as

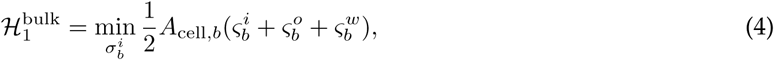
where the in-plane stresses of the IM and OM satisfy 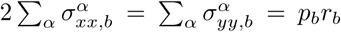 with *α* ∊ {*i, o, w*}, *p_b_* = *pV*_cell_/*V_b_*, where *V_b_* is the volume of the bulged cell, and *A*_cell,*b*_ = 2*πr_b_L_b_* − *A* is the surface area of the bulk. Here the subscript *_b_* denotes quantities of the bulged state. For any 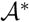, the minimizers of 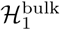 subject to the constraint 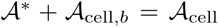 determine the dimensions of the bulk, where 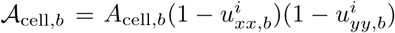. Given that the cylindrical bulk shrinks, and that the reference area by which it shrinks is relegated to the ellipsoidal bulge, the free energy of the bulged state can then be expressed as

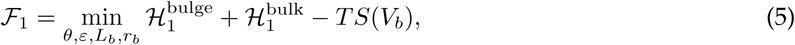
where the independent variables satisfy the reference area constraint. Here the aspect ratio *ε* = *b*/*a*, *S*(*V_b_*) is the entropy of mixing corresponding to a bulged volume 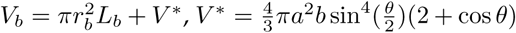 is the volume of the bulge, and the stretching energy of the bulge is

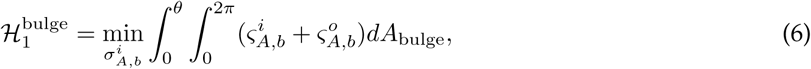
where, for *α* ∊ {*i, o*},

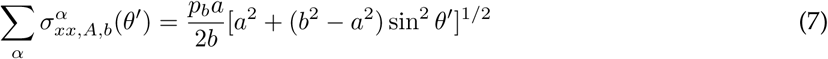
and

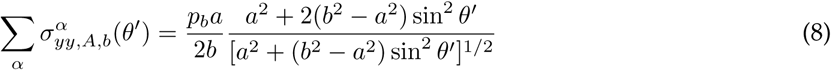

are the total in-plane stresses of the bulge as functions of parametric coordinates (*x*, *y*) = (*θ*′, *φ*). *θ*′ and *φ* denote the parametric angles along and around the axis of symmetry, respectively, and *dA*_bulge_ = *a* sin *θ*′[*a*^2^ + (*b*^2^ – *a*^2^) sin^2^ *θ*′]^1^/^2^*d φ dθ*′ is the surface area element of the bulge (SI).

Bulging is energetically favorable due to the relaxation of the entropic and stretching energies when 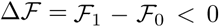. For characteristic parameter values relevant to *E. coli*, 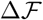 is calculated numerically as detailed in the SI and plotted in Fig. 2B. The individual contributions to 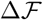 of both the changes in entropic energy and stretching energies of the bulk and the bulge are examined in the SI and Fig. S2. Although 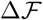 diverges as *θ* → 0—corresponding to the formation of a small bulge with diverging radii of curvature—there are alternate paths to the energetic minima for which no energetic barriers are present and bulging occurs spontaneously. One such path, in which a partial spherical bulge of fixed radius protrudes from the defect area before filling the defect area completely, is shown in Fig. 2C and analyzed in the SI. Importantly, for a wide range of *r_d_*, the configuration in which no bulging occurs is unstable, and the predicted bulge geometries appear consistent with experimental measurements (Fig. 3). For larger values of *r_d_*, bulges corresponding to smaller defects are, intriguingly, predicted to subtend larger angles, while the predictions for small *r_d_* can be supported by analytical calculations (SI). Moreover, our model predicts a volume increase on the order of 1-10% for characteristic values of *r_d_*, with larger volume changes corresponding to larger rd, and is consistent with a relaxation process in which the membranes slide against the wall due to the different strain rates of envelope components and the cell wall shrinks in the axial direction due to the membranes bearing larger loads (SI).

**Figure 3:**
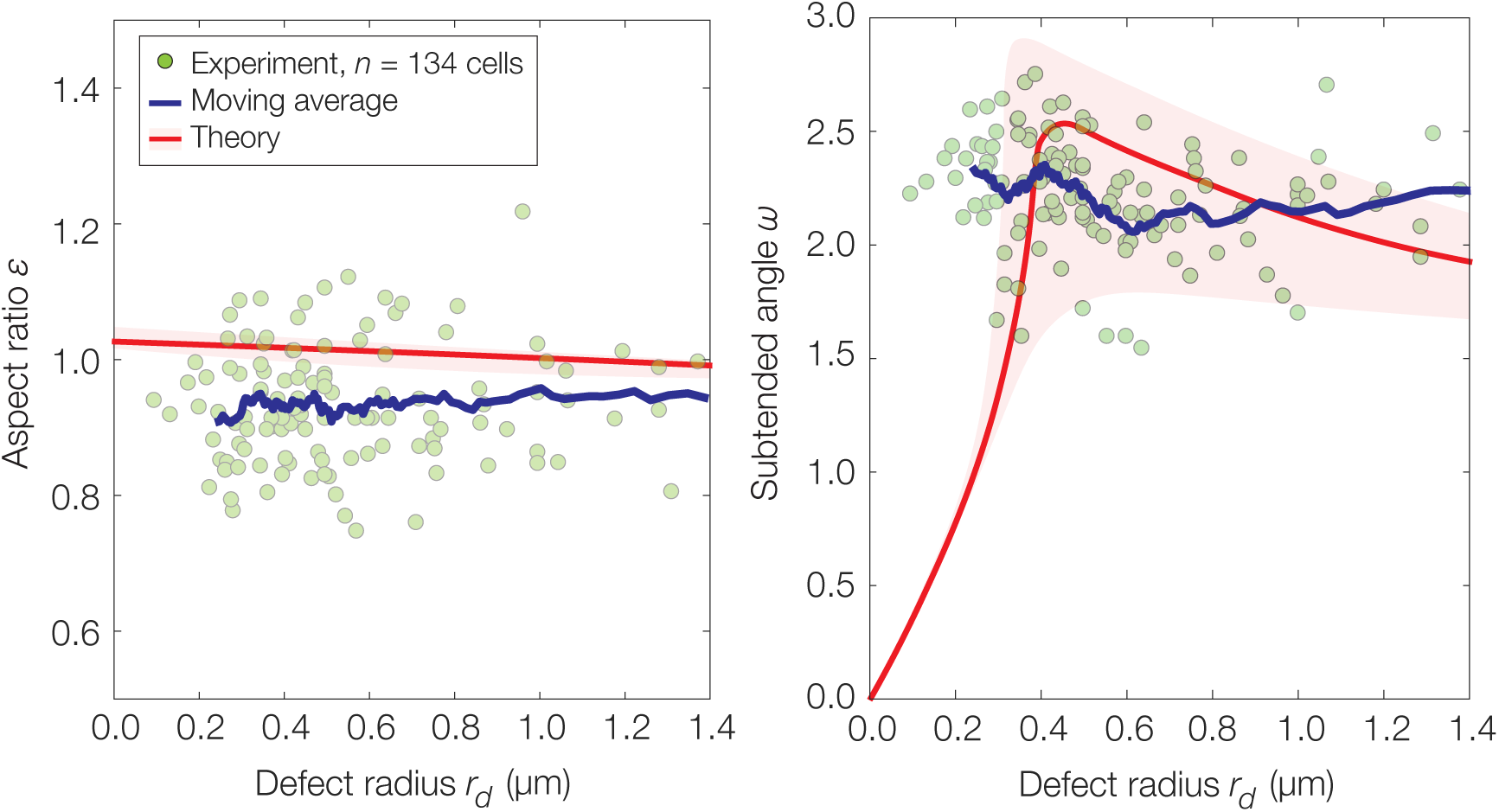
Statistics of bulged cells. Experimental measurements of bulge geometries for 134 cells. Here the aspect ratio is defined as *ε* = *b*/*a*. The curves represent moving averages and model predictions for the parameter values summarized in *Materials and Methods* and no fitting parameters. The shaded areas represent model predictions where the cell length is in the range [5 *µ*m,15 *µ*m], and the model predictions are less, or similarly, sensitive to similar changes in other parameters (not shown). The scatter indicates cell-to-cell variability.

We note three further implications of our analysis. First, bulging can be energetically favorable for a defect radius as small as *r_d_* = 10 nm (SI and Fig. S2). Second, over a large range of *r_d_*, the bulge volume *V*\* corresponding to minimizers of 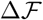 depends weakly on *L* (Fig. S3). The insensitivity to *L* suggests a physical explanation for the observation that cell length appears uncorrelated with bulge volume,^16^ and our model further predicts that bulge size does not significantly change over a broad range of *r*. Third, balancing the energy dissipation with the viscous drag on the bulge results in a timescale much smaller than 100 ms (SI), suggesting that the relaxation time may be limited by water flow: assuming the parameter values in *Materials and Methods*, the larger value of *L_p_* ≈ 10^−12^ m^3^/N · s implies a volumetric flow rate of approximately 20% of the initial cell volume per second and is consistent with a volume increase on the order of 1-10% of the initial cell volume during bulging. We anticipate further experiments, for instance ones which modulate membrane permeability during *β*-lactam killing, to elucidate the origins of these fast dynamics and test other predictions of our model.

### Model of swelling

During swelling, the amount of water uptake is determined by the same balance of the entropic and stretching energies of the cellular envelope as above: if lysis did not occur, then net flow into the cytoplasm would occur until the membranes are sufficiently stretched. In fact, the small synthesis rate of membrane material relative to water flow^49^ suggests that water flow is not limiting and that the membranes are always stretched. To support this notion, we analyzed the swelling of *E. coli* cells of different lengths over ~10 s and found that the population-averaged volumetric flow rate is small (Fig. 4A). In contrast, image analysis reveals that bulges grow at a rate consistent with theory when the defect radius, *r_d_*, also increases (Fig. 4B), implicating defect growth as the limiting step of bulge growth before lysis. Since the mean bulge radii at lysis are *a* ≈ 0.9 *µ*m and *b* ≈ 0.7 *µ*m, assuming the parameter values summarized in *Materials and Methods* and that turgor pressure has not changed due to osmotic stress responses^44,45,47,48,50,51,52^ suggests an upper bound for the yield areal strain of the *E. coli* IM and OM as approximately 20%, which lies within the empirical range of RBCs under impulsive stretching^53^ and exceeds that under quasi-static loading.^54^

**Figure 4:**
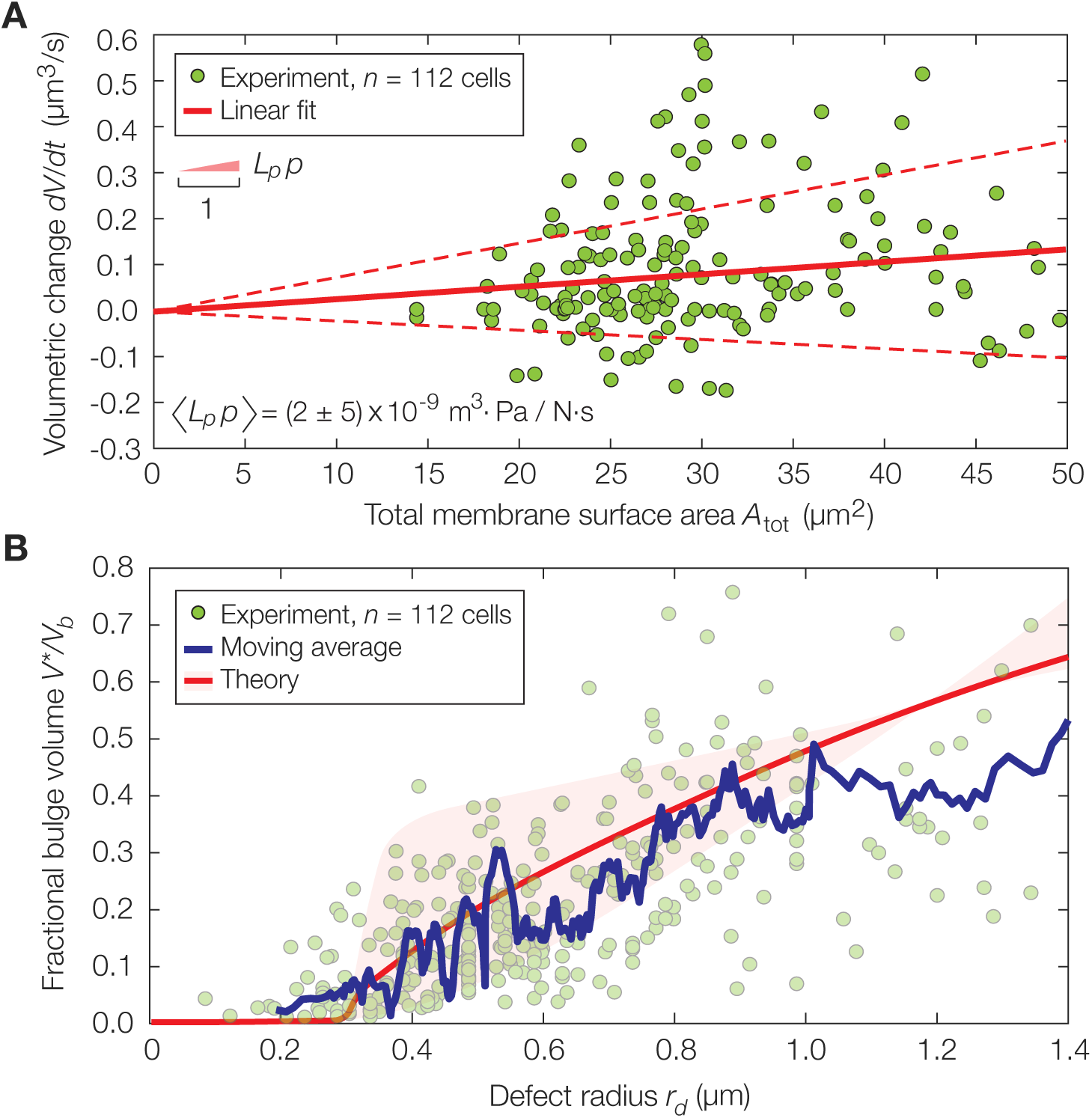
Statistics of swelling cells. (A) Plot of the volumetric flow rate *dV*/*dt* against the total membrane surface area *A*_tot_ for 112 swelling cells of different lengths and one or two data points per cell, with a linear fit overlaid. Error bars indicate one standard deviation (dashed lines). Here and in (B), the scatter indicates cell-to-cell variability. (B) Plot of *V*\*/*V_b_*, the fractional bulge volume, against the defect radius *r_d_* for the same cells as in (A), with the moving average and theoretical prediction overlaid. The parameter values are summarized in *Materials and Methods*. The shaded areas represent model predictions where the cell length is in the range [5 *µ*m,15 *µ*m], and the model predictions are less, or similarly, sensitive to similar changes in other parameters (not shown).

## Discussion

To summarize, we have used a continuum, elastic description of the cellular envelope to model membrane bulging and found that both continued water flow into the cytoplasm and defect enlargement can contribute to swelling. Our results underscore the different roles of each envelope component in resisting mechanical stresses and indicate that bulging can arise as a relaxation process mediated by membrane fluidity and water flow once a wall defect exists. These findings have broad implications on cellular physiology and morphogenesis. Because bulging and swelling result in eventual lysis and are mediated by cell wall defects, the existence of large pores in bacterial cell walls can be deadly. A growth mechanism which regulates pore size could help cells avoid lysis, in addition to regulating wall thickness and straight, rod-like morphology.^30^

Beyond bacterial morphogenesis, the combination of theory and experiment in our work has revealed a novel description of biological membrane physics and underscored the importance of mechanical stresses in cells. By being free to change their reference geometries, biological membranes differ from elastic shells, and we have shown that this difference has physiological implications on cell envelope mechanics and how mechanical stresses are distributed between membrane-solid layers. This paves the way for investigating and manipulating similar, rich interactions of fluid membranes with elastic surfaces^38^ and the material nature of living cells.

## Materials and Methods

### Model parameters

Unless otherwise specified, in this work we assume 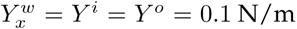, 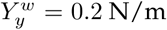, 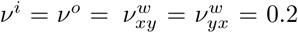, *p* = 0.5 atm, *T* = 300 K, *r* = 0.5 *µ*m, *L* = 10 *µ*m, and *r_d_*= 0.6 *µ*m. The curves in Fig. 2B are drawn with *L_b_* determined by the condition of fixed reference membrane area, *r_b_* = 0.5 *µ*m, and *ε* = 1, approximately the values at which the minimum of 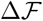 over *θ*, *r_b_*, and *ε* is achieved.

### Bacterial strains and growth

The wild-type strain used in this study is *E. coli* MG1655, and we verified that the morphological dynamics are statistically indistinguishable in two other wild-type strains, JOE309 and BW25113. Cells were grown in liquid LB (LB: 10g/L tryptone, 5g/L yeast extract, 10g/L NaCl) supplemented with no antibiotics. LB media containing 1.5% Difco agar (w/v) was used to grow individual colonies. Cells were taken from an overnight culture, diluted 100 to 1000-fold, and grown in LB at 37°C in a roller drum agitating at 60 rpm to an absorbance of approximately 0.3 to 0.6 (λ = 600 nm). Cells were then concentrated by centrifugation at 3000 rpm for 5 min and resuspended. We added 1 *µ*L of the bacterial culture to No. 1.5 coverslips (24×60 mm) and placed 1 mm thick LB agarose (1.5%) pads containing 50 *µ*g/mL of cephelaxin, a *β*-lactam antibiotic, on top for imaging. Cells were imaged immediately afterwards.

### Microscopy

We used a Nikon Ti inverted microscope (Nikon, Tokyo, Japan) equipped with a 6.5 *µ*m-pixel Hamamatsu CMOS camera (Hamamatsu, Hamamatsu City, Japan) and a Nikon 100x NA 1.45 objective (Nikon, Tokyo, Japan) for imaging. All cells were imaged at 37°C on a heated stage. The time between each frame during timelapse measurements ranged from 10 ms to 2 s, and the duration of timelapses varied from 10 min to 3 h. Images were recorded using NIS-Elements software (Nikon, Tokyo, Japan).

### Image analysis

Image sequences were compiled from previous work^16^ and from ten replicate experiments described above, which resulted in raw data for over 500 cells. These sequences were annotated manually in ImageJ (National Institutes of Health, Bethesda, MD) to obtain cell dimensions, bulge radii, defect radii, and subtended bulge angles. Bulged cells were fit to cylinders with protruding ellipsoids with radii (*a*, *a*, *b*), as described in the main text, to determine cell volumes and membrane areas. For Fig. 1D, a subset of 30 cells were chosen among cells which bulged on a timescale ~100 ms. Here and below, all cells considered bulged in the imaging plane. For Fig. 3, a subset of 134 cells were chosen among cells which bulged on a timescale ~1 s. For Fig. 4, a subset of 112 cells were chosen among cells which bulged on a timescale ~1 s and for which the cellular dimensions could be determined, and relevant statistics were measured or computed at two or three time points until ~10 seconds after bulging. This choice of timescale was made to mitigate the potential influence of cellular stress responses such as transport of solutes out of the cytoplasm,^44^ which could confound volumetric measurements. We discarded data points lying beyond the ranges plotted in Figs. 3 and 4, which corresponded to outliers, and applied a trailing moving average filter of 10 to 20 points to generate the moving average curves.

## Additional Information

Correspondence and requests for materials should be addressed to A.A.

## Acknowledgements

F.W. was supported by the National Science Foundation Graduate Research Fellowship under grant no. DGE1144152. A.A. was supported by the Materials Research and Engineering Center at Harvard, the Kavli Institute for Bionano Science and Technology at Harvard, the Alfred P. Sloan Foundation, and the Volkswagen Foundation. We thank John W. Hutchinson for numerous extended discussions, John W. Hutchinson, L. Mahadevan, Shmuel M. Rubinstein, Haim Diamant, Roy Kishony, Michael Moshe, Ugur Çetiner, and Zhizhong Yao for helpful feedback, Ethan C. Garner, Sean Wilson, and Georgia Squyres for microscopy assistance, Thomas G. Bernhardt and Sue Sim for the BW25113 strain, and Po-Yi Ho, Jie Lin, and Michael Moshe for comments on the manuscript.

## Author Contributions

F.W. and A.A. conceived the project, performed modeling, analyzed data, and wrote the paper. F.W. performed experiments.

## Supplementary Information

### Entropic origin of turgor pressure and stresses at equilibrium

Here we show that minimization of 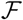 in equation (1) of the main text implies that, when the chemical potentials of water are equal both inside and outside the cell, the mechanical stresses in the cellular envelope are proportional to *p* = *kTC*, where *C* is the number density of solutes inside the cell (assuming no solutes outside the cell) and *p* is defined as the turgor pressure. For simplicity, we first neglect the differences between the cell wall, IM, and OM and consider the cellular envelope as a homogeneous, continuum, isotropic, elastic shell, with Young’s modulus and Poisson’s ratio *Y* and *ν*, respectively, and a fixed reference state. Similar conclusions can be shown to hold assuming orthotropic material properties over the entire cellular envelope and different geometries.

With notation similar to the main text, in this section we denote as *L*_1_ (*L*_0_) and *r*_1_ (*r*_0_) the (reference) length and radius, respectively, of the cell envelope. Assuming the linear strain-displacement relations in the main text and setting

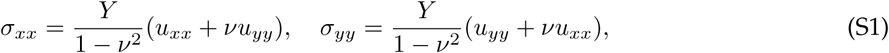

the stretching energy is

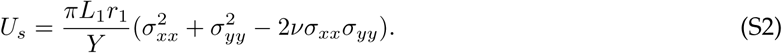

We ignore the bending energy, as discussed below, and consider now the entropic term. We assume the dilute limit of a small, fixed number of solutes *n_s_* inside the cell, so that *n_s_*/*n_w_* ≪ 1, where *n_w_* denotes the number of water molecules. In this case, and assuming that the volume occupied per solute molecule is comparable to that occupied per water molecule, the number of water molecules contained in the cell envelope can be approximated as 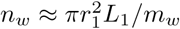, where *m_w_* ≈ 30 × 10^−30^ m^3^ is the volume occupied per water molecule. By substituting these expressions into the entropy of mixing discussed in the main text, we find that the entropic term is approximately

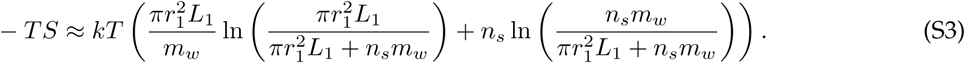

Since the cell envelope geometry may change, we minimize 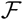 with respect to the stresses *σ_xx_* and *σ_yy_*. Anticipating that *L*_1_ = *L*_0_ + *δL* and *r*_1_ = *r*_0_ + *δr* and that *u_xx_* = *δL*/*L*_0_, *u_yy_* = *δr*/*r*_0_, and *n_s_*/(*n_s_* + *n_w_*) are small, expanding 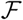 to second order in both *u_xx_* and *u_yy_* and first order in *n_s_*/(*n_s_* + *n_w_*) around zero and solving for the values of *δL* and *δr* which minimize 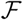 yield

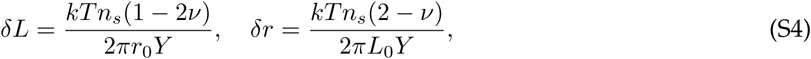

which implies

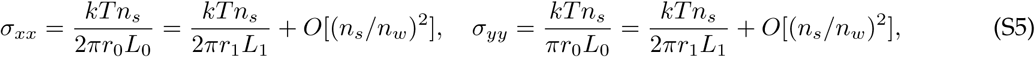

consistent with loading by a pressure *p* = *kTC*, where 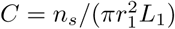. From this, we also find that *n_s_* can be expressed in terms of cellular parameters as 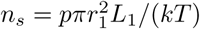.

### Bending energies are negligible even at the neck

Throughout this work, we have assumed that the bending energies are negligible compared to the stretching energies. The bending energy of an isotropic shell is *E*_bend_ = 2*k_b_* ∫ *H*^2^*dA*. Here *k_b_* is the bending rigidity, *H* is the mean curvature, a vanishing spontaneous curvature is assumed for all surfaces for simplicity, and the contribution of Gaussian curvature to the elastic energy is ignored due to the Gauss-Bonnet theorem and absence of topological change. The bending energy 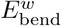 of the orthotropic cell wall assumes a more complicated form involving bending rigidities in the *xx*, *xy*, and *yy* directions [1]. However, here we do not consider bending deformations of the cell wall. We therefore leave the form of 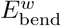 unspecified and ignore it in the following. We now consider the addition of these bending energies to the analysis in this study. The combined bending energy of the unbulged state, a cylinder, is 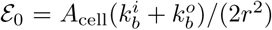, where we drop the subscripts *_u_* and _*b*_ on *r* for simplicity and retain the notation used in the main text. The combined bending energy of the bulged state is

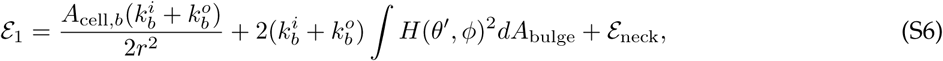
where we compute the mean curvature of an ellipsoid parameterized by (*α* sin *θ*′ cos *ϕ*, *a* sin *θ*′ sin *ϕ*, *b* cos *θ*′) from its second fundamental form as

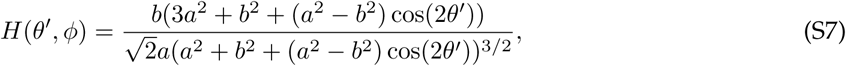
and 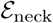 is the bending energy of the neck. For an ellipsoidal bulge joined to a cylinder, the mean curvature diverges at the kink of the neck. In lieu of a perfect kink, we may suppose instead that the geometry of the neck is described by a partial, circular torus of major and minor radii *D* and *C* (Fig. S1). *C* can be set to satisfy conservation of membrane reference surface areas, so that the reference area of the torus is identical to the reference area of the neck that it replaces; however, *D* is constrained by the bulge radius to be *D* ≈ *a* sin *θ*. For the case in which the parametric angle *θ* ≤ *π*/2, the sector of the toroidal cross-section needed to bridge the neck can be taken to be < *θ* with its value dependent on the choice of *C*, while for *θ* > *π* /2, half of the cross-section suffices over a range of *C* (Fig. S1). Then, as the area of the toroidal neck is bounded by 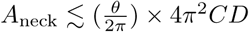, the bending energy of the toroidal neck satisfies

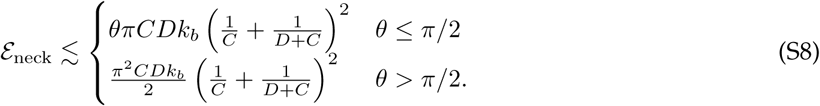

Introducing a toroidal neck also results in a contribution to the stretching energy. The in-plane stress for a circular torus are [2, 3]

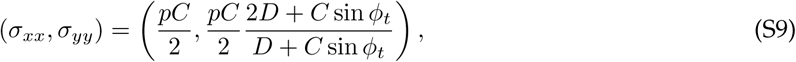
where *θ*_*t*_ and *ϕ*_*t*_ are respectively the toroidal and poloidal angles of the torus and (*x*, *y*) = (*θ*_*t*_, *ϕ*_*t*_); as the typical volume changes we consider are small, for simplicity we ignore the effect of solute dilution on p here and in the remainder of the SI, unless specified otherwise, and note that this does not significantly change our results. Taking 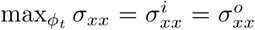 and 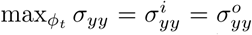, we obtain an additional stretching energy of the neck upper bounded by 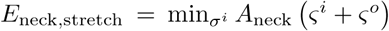. Finally, introducing the neck increases the volume enclosed by the IM, and considering its entropic contribution explicitly would only decrease the free energy change further. Thus, the free energy change 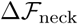 due to bulging in the case where a kink at the neck is replaced by a partial torus is upper bounded by

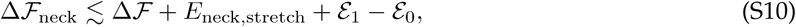
where 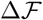 is of the same form as that considered in the main text. For the parameter values indicated in *Materials and Methods* and *k_b_* = 20 *kT*, *D* ~ 1 *µ*m, and *C* ~ 10 nm, we find that the corrections indicated in equation (S10) are of lower orders of magnitude (~ 10^−16^ to 10^−17^ J) than the energy scales considered in the main text and do not change our results; neither do they present energetic barriers to relaxation. Taken together, these considerations suggest that it is indeed justifiable to neglect the bending energies and the energetic contribution of the neck.

### Stress analysis of an ellipsoidal bulge and calculation of undeformed surface area

We consider the ellipsoidal surface parameterized by (*a* sin *θ*′ cos *ϕ*, *a* sin *θ*′ sin *ϕ*, *b* cos *θ*′). The stress resultants of the bulge are found by solving the membrane equations

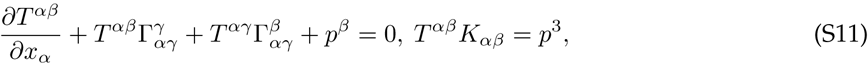
where *T* is the stress resultant tensor, *K* is the curvature tensor, 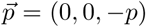 is the external force per unit area, 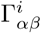 are Christoffel symbols of the second kind, and the indices (*α*, *β*, *γ*) range over *θ*′ ≡ *x* ≡ *x*_1_ and *ϕ* ≡ *y* ≡ *x*_2_ [4, 5]. (Here ^3^ denotes surface normal quantities.) The in-plane stresses correspond to the stress resultant tensor with mixed (raised and lowered) indices. Upon index lowering by the covariant metric tensor *G_αβ_* of the deformed state, where

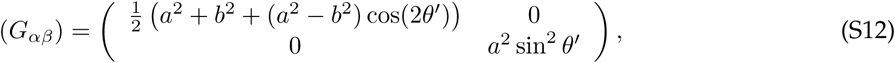

these give, as functions of the parametric angle along the axis of symmetry *θ*′,

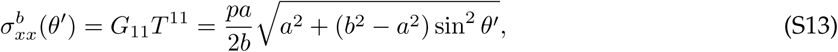

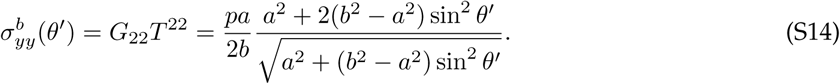

Note that, assuming the linear strain-displacement relation *E_αβ_* = *e_αβ_* = (*G_αβ_* − *g_αβ_*)/2 for a thin shell, where *E* (*e*) is the strain tensor of the deformed (reference) state of the bulge, index lowering with the covariant metric tensor of the undeformed state, *g_αβ_*, and retaining the terms accurate to linear order in the strains results in identical expressions. As we employ a linear theory in this work, we shall continue to lower indices with *G_αβ_* below.

In the main text, we wish to determine 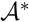, the reference surface area of an ellipsoidal bulge. Although the linearized strain-displacement relations of a thin shell are analytically and numerically difficult to solve for the ellipsoidal geometry we consider, 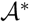 can be calculated directly from the metric tensor using the strain-displacement relation *E_αβ_* = (*G_αβ_* − *g_αβ_*)/2. In particular, we set 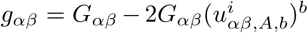, where 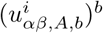 is determined from equations (7) and (8) of the main text and the constitutive relation, and take 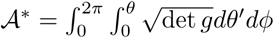.

### Numerical minimization of 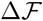

As the analytical calculations below suggest, the minimization of 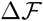 is algebraically and analytically complex. We numerically minimized 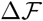 by determining 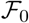 and then determining 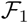. To find the minimizers corresponding to equation (3) of the main text, we first discretized *r_u_* over the interval [0.3 *µ*m, 0.7 *µ*m] into 4 steps. As the IM and OM share identical material properties for the parameter values considered in this work, by symmetry we set 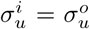 and 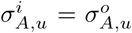, and likewise for determining 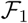 below. Hence, for a given a value of *r_u_*, *p_u_* and *L_u_* can be determined self-consistently by the membrane reference surface area constraint, and a numerical value of the expression in equation (3) of the main text can be computed. We took 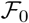 to be the minimal value obtained by sampling over *r_b_* in this way.

To determine the minimizers corresponding to 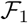 in equation (5) of the main text, we discretized *dr* = *r* − *r_b_* over the interval [−0.2 *µ*m, 0.2 *µ*m] into 4 steps, *ε* over the interval [0.8,1.2] into 10 steps, *θ* over the interval [0, *π*] into 200 steps, and determined *p_b_* and *L_b_* self-consistently from the membrane reference surface area constraint. For given values of each sampled variable, a numerical value of 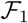 can therefore be computed. We omitted computations corresponding to cases where *L_b_* < 2*r_d_*, in which case the geometry assumed in our model is unphysical. For Fig. 3 in the main text, we determined the model predictions as follows: for given values of *r_b_* and *ε*, we considered the first local minimum, 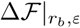, of 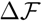 viewed as a function of *θ* starting from *θ* = 0. We then took the predicted configuration to correspond to the minimum of 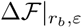 over all sampled values of *r_b_* and *ε*.

The foregoing calculations were repeated for variations in *r_d_*, *r*, and *L*. We discretized *r_d_* over the interval [0 *µ*m,1.4 *µ*m] into 28 steps, *r* over the interval [0.3 *µ*m,1.1 *µ*m] into 8 steps, and *L* over the interval [2 *µ*m, 20 *µ*m] into 18 steps, interpolated the model predictions, and smoothed the resultant curves. We repeated the foregoing calculations with different discretizations and ranges and verified that our model predictions were not significantly changed.

### Changes in the entropic and stretching energies

In this section, we consider the changes in entropic energy, stretching energy of the ellipsoidal bulge, and stretching energy of the cylindrical bulk separately, which we respectively denote as Δ*TS*, 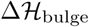, and 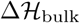. 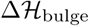 and 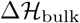 are the differences of the first and second terms of equations (4) and (6) of the main text, respectively. For the parameter values assumed in *Materials and Methods*, plots of Δ*TS*, 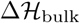, 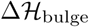, 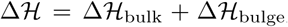, and 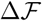 as functions of *V*\*/*V_b_* are shown in Fig. S2A. We note, in particular, that the trade-off between the stretching and entropic energies determines bulge size, and that both bulge growth and shrinking of the cylindrical bulk yield similar contributions to the increase in stretching energy in the energy-minimizing conformation.

### Bulging occurs spontaneously

Here we demonstrate the existence of a path to the energy-minimizing state shown in Fig. 2B of the main text which is monotonically decreasing in the free energy. In particular, this path avoids the divergence of 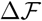 as *θ*, *V\** → 0 shown in Fig. 2B of the main text. We consider extruding a sphere of a fixed radius *R* ≥ *r_d_* through the wall defect until the cross-section of the sphere coincides with the wall defect. Reusing notation introduced elsewhere in the text, the free energy of the cell envelope with a spherical bulge of fixed radius *R*, but varying parametric angle *θ*, protruding through a defect can be written as

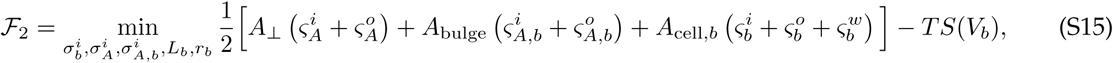
where *A*_⊥_ = *A* − *A_e_*, 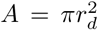 is the defect area, 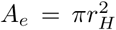 is the cross-sectional bulge area with *r_H_* = *R* sin *θ* ≤ *r_d_*, *A*_bulge_ = 2*πR*^2^ (1 – cos *θ*), *A*_cell,*b*_ = 2*πr_b_L_b_* − *A* is the surface area of the remaining cylinder, and *S*(*V_b_*) is the entropy of mixing corresponding to a total volume 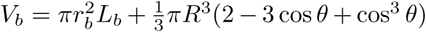. The reference area constraint can be written as 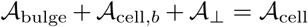, where the undeformed surface areas can be determined from the deformed surface areas in a straightforward manner similar to that considered in the main text. The conditions of force balance on each region are 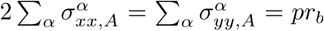, 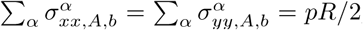, and 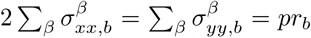, where *α* ∈ {*i*, *o*} and *β* ∈ {*i*, *o*, *w*}. For the parameter values considered in *Materials and Methods*, a plot of 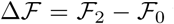 which is monotonically decreasing in *θ* until *θ* = *π*/2 is shown in Fig. S2B.

### Analytical calculations under simplified model assumptions

Here we consider analytical calculations in order to support the numerical results presented in the main text. Due to the complicated form of 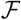, analytical calculations may only be tractable in certain limits and under simplifying assumptions. In this section, in order to simplify analytical calculations and provide further intuition for the results presented in the main text, we neglect the existence of the outer membrane and assume that 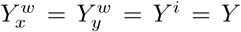, 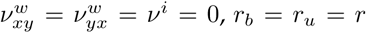, and *ε* = 1. We consider the dependence of the subtended angle on the defect radius in the limit where the defect radius is small.

Under the assumptions above, it is possible to solve for 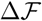 analytically. Doing so results in algebraically complex forms for 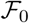 and 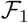:

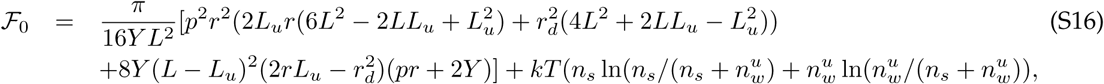
where *L_u_* is constrained by conservation of membrane reference surface area to yield

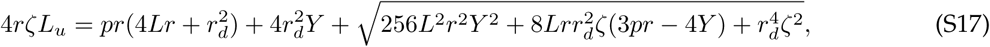
where *ζ* = *pr* + 4*Y*, *n_s_* = *pπr*^2^*L*/(*kT*), and 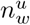, the number of water molecules in the unbulged state, is 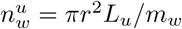, and

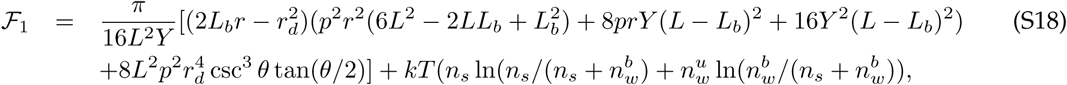

Where

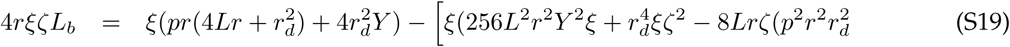

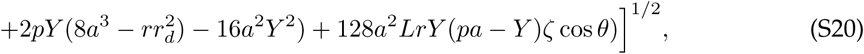

*ξ* = *pr* − 2*Y*, *a* = *r_d_*/sin *θ*, and 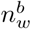, the number of water molecules in the bulged state, is 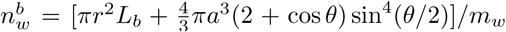. Based on our numerical results, we anticipate that *r_d_* ~ *θ* in the limit of small *θ*. Expanding 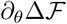 to fourth order in *r_d_*/*r* around 0 and first order in *θ* around 0 and solving for its nontrivial zeros then gives the following relation between *r_d_* and *θ*:

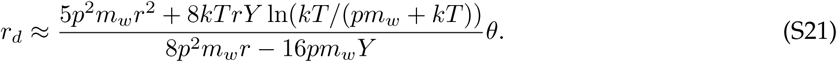

Equation (S21) describes a model prediction for the subtended (parametric) angle, *θ*, as a function of the defect radius, *r_d_*, under the simplifying assumptions discussed above. It predicts that, for small bulges, the subtended angle is characteristically small and increases with the defect radius, consistent with the numerical results presented in Fig. 3 of the main text and Fig. S3. For the parameter values considered in this work and *Y* = 0.1 N/m, the prefactor of equation (S21) is approximately 0. 24 *µ*m and the prediction of equation (S21) is consistent with Fig. 3 of the main text.

### Variables of the bulged state

For the parameter values considered in *Materials and Methods*, the variables of the bulged state are *a* ≈ 0.8 *µ*m, *ε* ≈ 1, *θ* ≈ 2.3, *L*_1_ ≈ 8.8 *µ*m, and *r*_1_ ≈ 0.5 *µ*m, while the fractional volume increase of the entire cell is Δ*V* ≈ 0.1. These variables attain similar values over a large range of *r_d_*, from *r_d_* = 0.2 *µ*m to *r_d_* = 1 *µ*m; notably, Δ*V* ranges from approximately 0 to 0.5. Our model predicts that, due to the different strain rates between envelope components, the membranes may slide against the cell wall as the cell bulges. Indeed, for the parameter values considered in *Materials and Methods*, the membrane strains in the bulk are identical between the IM and OM and change from 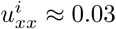 and 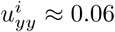 to 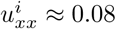 and 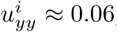, while the wall strains change from 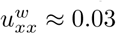 and 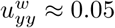 to 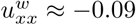 and 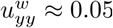. Intriguingly, our model predicts that for typical parameter values the cell wall is compressed axially. In this case, it is energetically favorable for membrane area to be appropriated by shortening the bulk beyond the reference length of the cell wall.

### Critical defect radius for bulging

In previous work [6], a critical defect radius of *r_d_* ≈ 20 nm for bulging was found by considering the trade-off between the bending and pressure-volume energies. The bending energy of a hemispherical bulge of radius *r_d_* is *E*_bend_ = 4*πk_b_*, while the pressure-volume energy is 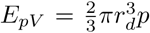. For the parameter values considered in this work, neglecting the bending energy at the bulge neck, and considering both the IM and OM in the presence of sufficient excess area, 2*E*_bend_ = *E_pV_* when *r_d_* = 21 nm, consistent with the estimate in [6]. However, allowing for membrane reorganization, a critical defect radius for bulging may exist by considering primarily the trade-off between the bending and stretching energies. In this case, the decrease in free energy can be caused primarily by a decrease in the stretching energy, which is dependent on the shape of the cell envelope: the stretching energy may be lowered if the bulge, for which the stresses are on the order of ~ *pr_d_*, replaced the cylindrical geometry, for which the stresses are proportional to *pr*. The stretching energy saved by bulging is well approximated by the contribution over the defect, 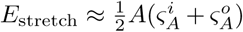. For the parameter values considered in this work, 2*E*_bend_ < *E*_stretch_ even when *r_d_* ≈ 10 nm, approximately the thickness of a lipid bilayer. To support this estimate, Fig. S2C shows a plot of 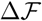 for *r_d_* = 10 nm and the parameter values considered in *Materials and Methods*.

### Timescale of the bulging response against viscous drag

We show that balancing the energy change computed above with the viscous drag on the bulge implies a timescale that is smaller than 100 ms, and hence energy dissipation cannot account for the observed timescale of bulging. The characteristic scale of 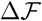 in our work is 10^−14^ J, while the power dissipation due to viscous drag on an expanding sphere is 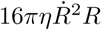, where *η* denotes the medium viscosity and *R* is the radius of the sphere [7]. Supposing the viscosity of water, *η* = 10^−3^ Pa · s, and estimating *R* = 0.5 *µ*m and 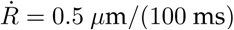 = 0.5 *µ*m/(100 ms) then results in an energy scale of 10^−19^ J/s. Equivalently, a power dissipation of 10^−14^ J/(100 ms) implies a bulging timescale of ~ 0.1 ms.

## Supplementary Videos

Supplementary Video 1: **Lysis dynamics of *E. coli* cells.** Supplementary Video 1 shows a population of wild-type *E. coli* cells bulging, swelling, and lysing under antibiotic treatment. The time between frames is 30 seconds, the timelapse covers a period of approximately 1 hour, and the field of view is 100 *µ*m × 80 *µ*m.

## Supplementary Figures

**Figure S1:**
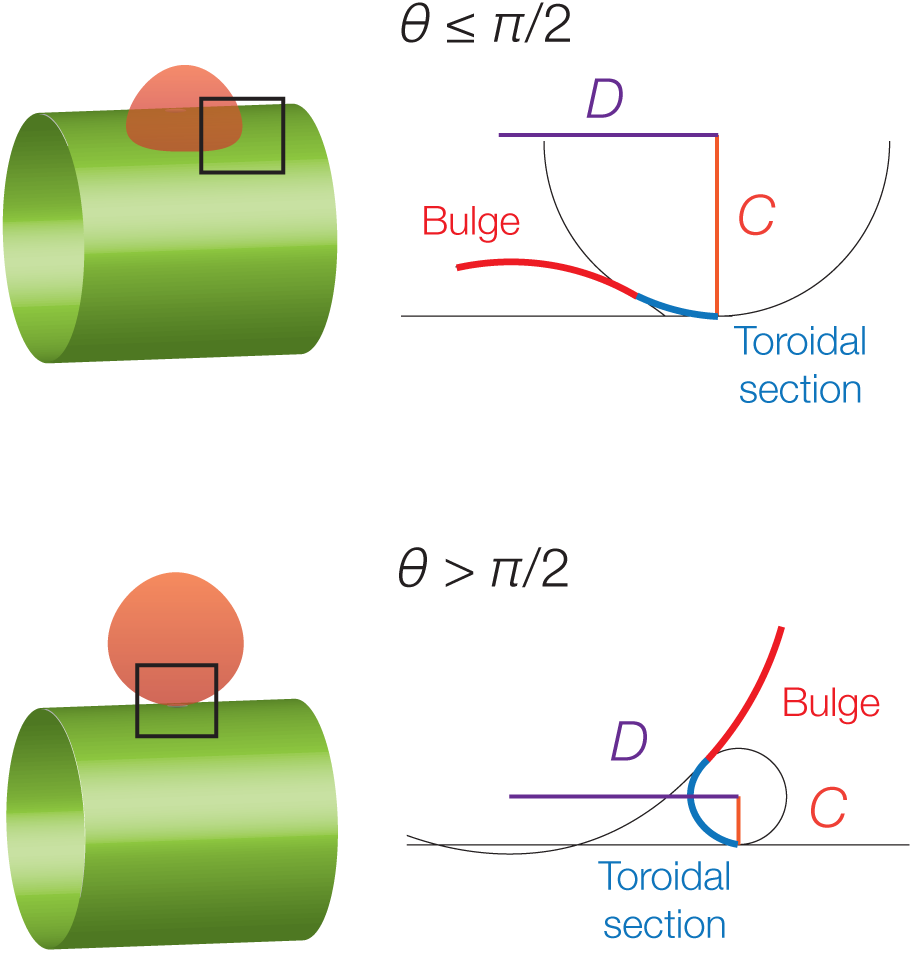
Bending energy at the neck. To avoid the divergence of the bending energy at the neck, we connect the bulge to the cylindrical bulk with a partial torus and estimate the resulting energetic contribution. The planar diagrams show cross-sections of the axisymmetric geometries.

**Figure S2:**
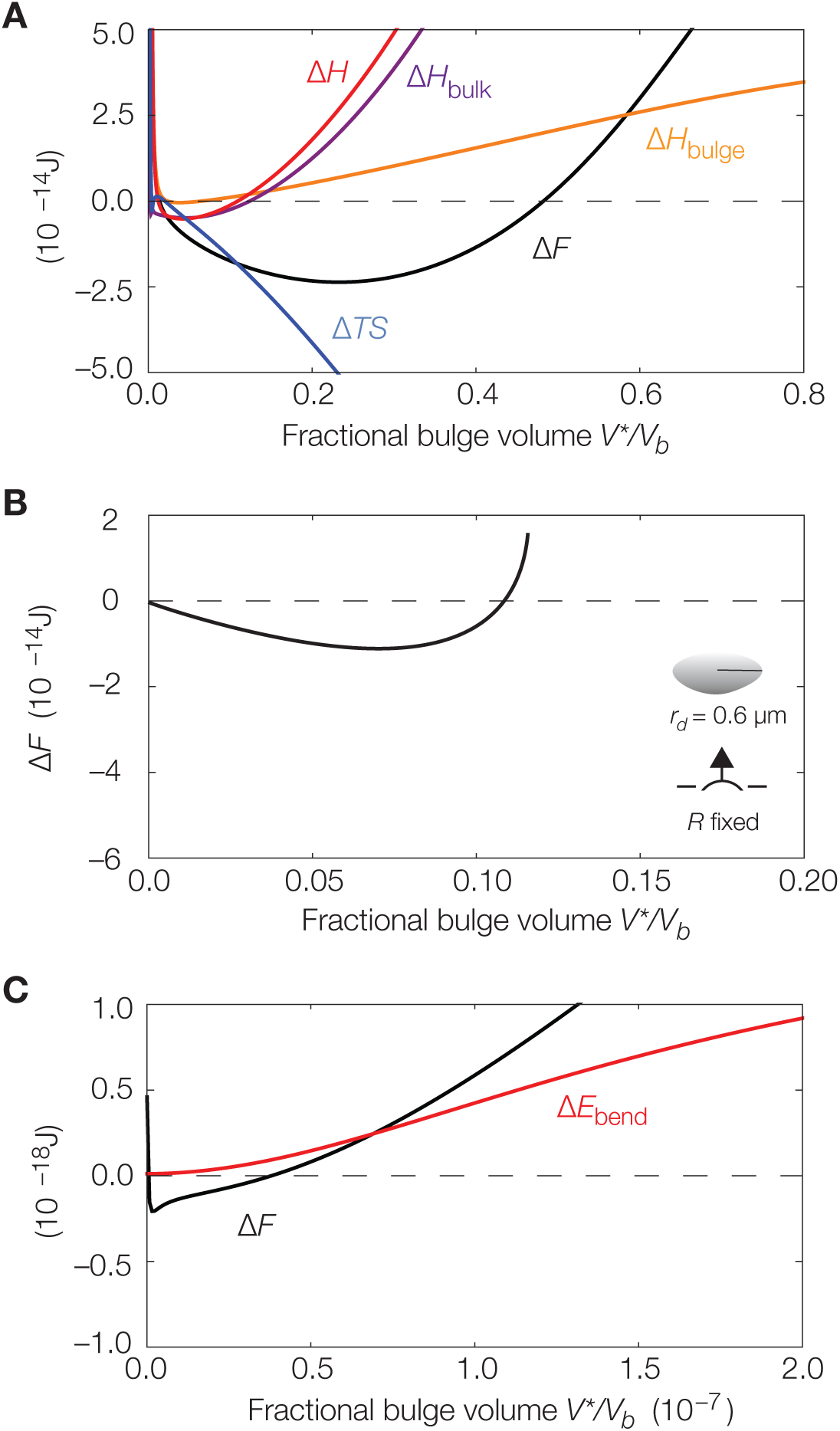
Energetics of bulging. (A) Contributions to the free energy change: same as Fig. 2B in the main text, but with individual contributions to 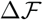 shown. (B) Bulging can occur spontaneously: there are paths that are monotonically decreasing in free energy to the energetic minimum shown in Fig. 2B of the main text. For the same parameter values as in *Materials and Methods*, a plot of the free energy change due to the extrusion of a spherical bulge with a fixed radius *R* = 0.6 *µ*m is shown. The curve is truncated when the subtended angle is *π*. (C) Bulging is energetically favorable even for small defects: same as Fig. 2B in the main text, but with *r_d_* = 10 nm and the energy change Δ*E*_bend_ = 2*k_b_A*_bulge_/*a*^2^, where *A*_bulge_ is the area of the bulge, corresponding to bending of the bulge (red curve) and the free energy change neglecting the bending energies (black curve) shown.

**Figure S3:**
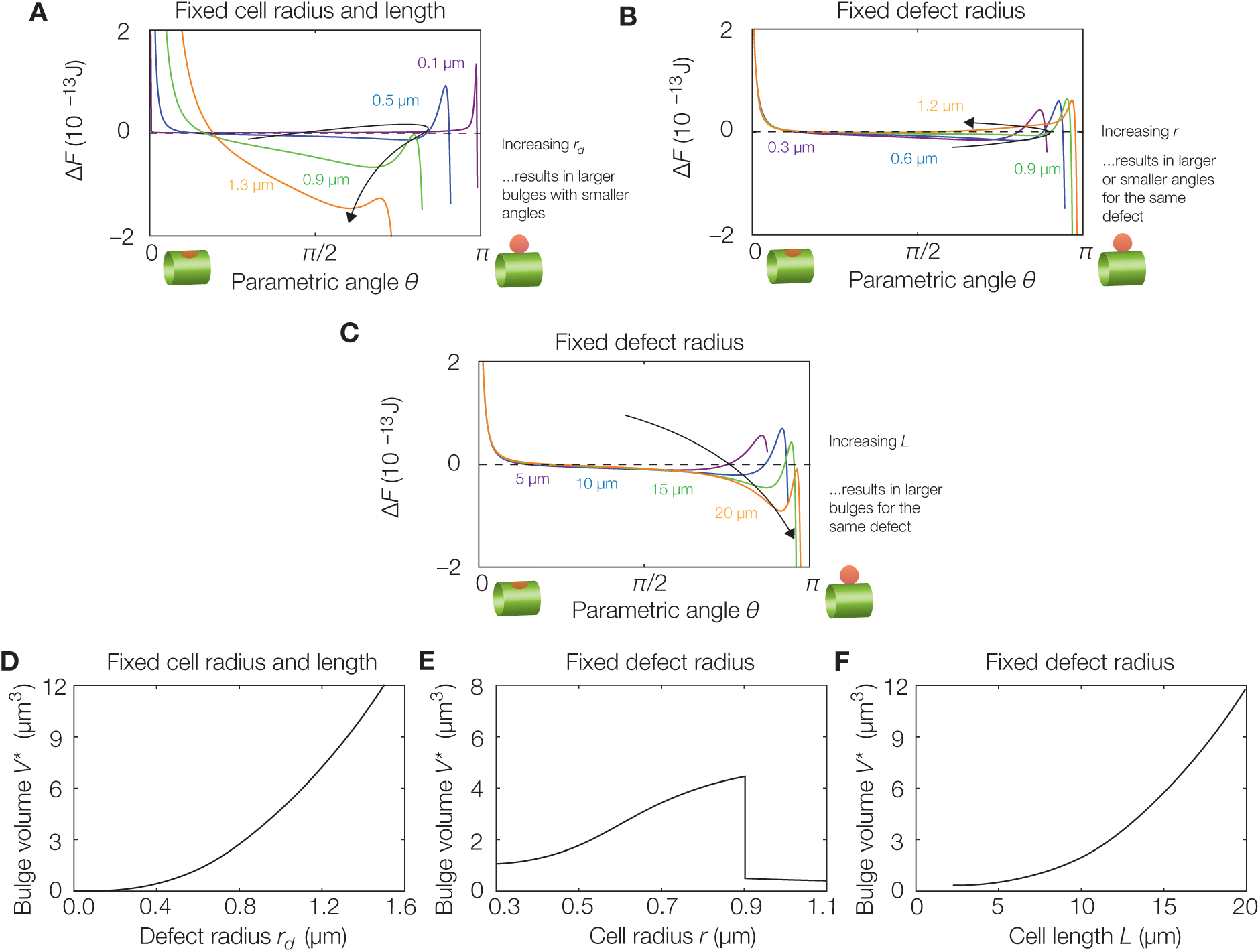
Dependence of bulge size on cell length, cell radius, and defect radius. (A) Plot of 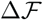 against the parametric angle *θ* for the parameter values summarized in *Materials and Methods* and varying *r_d_*. For simplicity, all curves are drawn assuming *r_b_* = *r*, *ε* = 1, and *L_b_* determined by the reference area constraint, but the plots are qualitatively similar over a wide range of parameter values. The curves are truncated when *L_b_* < 2*r_d_*, in which case the geometry assumed in our model is unphysical. (B) Same as (A), but with varying *r*. (C) Same as (A), but with varying *L*. (D) Predicted dependence of bulge volume on defect radius, assuming the parameters summarized in *Materials and Methods*. (E) Same as (D), but for the dependence of bulge volume on cell radius. The non-monotonicity is consistent with the transition between the regimes of large and small subtended angles shown in Fig. 3 of the main text. (F) Same as (D), but for the dependence of bulge volume on cell length.

## References

1. World Health Organization. Antimicrobial resistance: global report on surveillance (WHO, 2014).

2. Ventola, C. L. The antibiotic resistance crisis, part 1: causes and threats. P. T. 40, 277–283 (2015).

3. Bao, G. & Suresh, S. Cell and molecular mechanics of biological materials. Nat. Mater. 2, 715–725 (2003).

4. Guillot, C. & Lecuit, T. Mechanics of epithelial tissue homeostasis and morphogenesis. Science 340, 1185–1189 (2013).

5. Ladoux, B. & Mège, R.-M. Mechanobiology of collective cell behaviours. Nat. Rev. Mol. Cell Biol. 18, 743–757 (2017).

6. Lederberg, J. Mechanism of action of penicillin. J. Bacteriol. 73, 144 (1957).

7. Hahn, F. E. & Ciak, J. Penicillin-induced lysis of *Escherichia coli*. Science 125, 119–120 (1957).

8. Benveniste, R. & Davies, J. Mechanisms of antibiotic resistance in bacteria. Annu. Rev. Biochem. 42, 471–506 (1973).

9. Tomasz, A. The mechanism of the irreversible antimicrobial effects of penicillins: how the beta-lactam antibiotics kill and lyse bacteria. Annu. Rev. Microbiol. 33, 113–137 (1979).

10. Kohanski, M. A., Dwyer, D. J., Hayete, B., Lawrence, C. A. & Collins, J. J. A common mechanism of cellular death induced by bactericidal antibiotics. Cell 130, 797–810 (2007).

11. Kohanski, M. A., Dwyer, D. J. & Collins, J. J. How antibiotics kill bacteria: from targets to networks. Nat. Rev. Microbiol. 8, 423–435 (2010).

12. Cho, H., Uehara, T. & Bernhardt, T. G. Beta-lactam antibiotics induce a lethal malfunctioning of the bacterial cell wall synthesis machinery. Cell 159, 1300–1311 (2014).

13. Höltje, R. V. Growth of the stress-bearing and shape-maintaining murein sacculus of *Escherichia coli*. Microbiol. Mol. Biol. Rev. 62, 181–220 (1998).

14. Amir, A. & van Teeffelen, S. Getting into shape: How do rod-like bacteria control their geometry? Syst. Synth. Biol. 8, 227–235 (2014).

15. Young, K. D. The selective value of bacterial shape. Microbiol. Mol. Biol. Rev. 70, 660–703 (2006).

16. Yao, Z., Kahne, D. & Kishony, R. Distinct single-cell morphological dynamics under beta-lactam antibiotics. Mol. Cell 48, 705–712 (2012).

17. Koch, A. L. Bacterial Growth and Form (Springer Science & Business Media, 2001).

18. Chung, H. S. et al. Rapid beta-lactam-induced lysis requires successful assembly of the cell division machinery. Proc. Natl. Acad. Sci. USA 106, 21872–21877 (2009).

19. Huang, K. C., Mukhopadhyay, R., Wen, B., Gitai, Z. & Wingreen, N. S. Cell shape and cell wall organization in Gram-negative bacteria. Proc. Natl. Acad. Sci. USA 105, 19282–19287 (2008).

20. Din, M. O. et al. Synchronized cycles of bacterial lysis for *in vivo* delivery. Nature 536, 81–85 (2016).

21. Lee, A. J. et al. Robust, linear correlations between growth rates and *β*-lactam-mediated lysis rates. Proc. Natl. Acad. Sci. USA 115, 4069–4074 (2018).

22. Charras, G. T., Yarrow, J. C., Horton, M. A., Mahadevan, L. & Mitchison, T. J. Non-equilibration of hydrostatic pressure in blebbing cells. Nature 435, 365–369 (2005).

23. Charras, G. T., Coughlin, M., Mitchison, T. J. & Mahadevan, L. Life and times of a cellular bleb. Biophys. J. 94, 1836–1853 (2008).

24. Lim, F. Y., Chiam, K. H. & Mahadevan, L. The size, shape, and dynamics of cellular blebs. EPL 100, 28004 (2012).

25. Charras, G. T. A short history of blebbing. J. Microscopy 231, 466–478 (2008).

26. Fackler, O. T. & Grosse, R. Cell motility through plasma membrane blebbing. J. Cell Biol. 181, 879–884 (2008).

27. Charras, G. T., Hu, C.-K., Coughlin, M. & Mitchison, T. J. Reassembly of contractile actin cortex in cell blebs. J. Cell Biol. 175, 477–490 (2006).

28. Daly, K. E., Huang, K. C., Wingreen, N. S. & Mukhopadhyay, R. Mechanics of membrane bulging during cell-wall disruption in Gram-negative bacteria. Phys. Rev. E Stat. Nonlin. Soft Matter Phys. 83, 041922 (2011).

29. Hussain, S. et al. MreB filaments align along greatest principal membrane curvature to orient cell wall synthesis. eLife 7, e32471 (2018).

30. Wong, F. et al. Mechanical strain-sensing implicated in cell shape recovery in *Escherichia coli*. Nat. Microbiol. 2, 17115 (2017).

31. Calladine, C. R. Theory of Shell Structures (Cambridge University Press, 1983).

32. Santangelo, C. D. Buckling thin disks and ribbons with non-Euclidean metrics. EPL 86, 34003 (2009).

33. Deng, Y., Sun, M. & Shaevitz, J. W. Direct measurement of cell wall stress stiffening and turgor pressure in live bacterial cells. Phys. Rev. Lett. 107, 158101 (2011).

34. Lan, G., Wolgemuth, C. W. & Sun, S. X. Z-ring force and cell shape during division in rod-like bacteria. Proc. Natl. Acad. Sci. USA 104, 16110–16115 (2007).

35. Sun, Y., Sun, T.-L. & Huang, H. W. Physical properties of *Escherichia coli* spheroplast membranes. Biophys. J. 107, 2082–2090 (2014).

36. Waugh, R. & Evans, E. A. Thermoelasticity of red blood cell membrane. Biophys. J. 26, 115–131 (1979).

37. Phillips, R., Kondev, J., Theriot, J. & Garcia, H. Physical Biology of the Cell (Garland Science, 2012).

38. Staykova, M., Holmes, D. P., Read, C. & Stone, H. A. Mechanics of surface area regulation in cells examined with confined lipid membranes. Proc. Natl. Acad. Sci. USA 108, 9084–9088 (2011).

39. Amir, A., Babaeipour, F., McIntosh, D. B., Nelson, D. R. & Jun, S. Bending forces plastically deform growing bacterial cell walls. Proc. Natl. Acad. Sci. USA 111, 5778–5783 (2014).

40. Tuson, H. H. et al. Measuring the stiffness of bacterial cells from growth rates in hydrogels of tunable elasticity. Mol. Microbiol. 85, 874–891 (2012).

41. Jadidi, T., Seyyed-Allaei, H., Tabar, M. R. R. & Mashagh, A. Poisson’s ratio and Young’s modulus of lipid bilayers in different phases. Front. Bioeng. Biotechnol. 2, 8 (2014).

42. Yao, X., Jericho, M., Pink, D. & Beveridge, T. Thickness and elasticity of Gram-negative murein sacculi measured by atomic force microscopy. J. Bacteriol. 181, 6865–6875 (1999).

43. Sperelakis, N. Cell Physiology Source Book: Essentials of Membrane Biophysics (Academic Press, 1995).

44. Buda, R. et al. Dynamics of *Escherichia coli*’s passive response to a sudden decrease in external osmolarity. Proc. Natl. Acad. Sci. USA 113, E5838–E5846 (2016).

45. Çetiner, U. et al. Tension-activated channels in the mechanism of osmotic fitness in *Pseudomonas aeruginosa*. J. Gen. Physiol. 149, 595–609 (2017).

46. Peterlin, P., Arrigler, V., Haleva, E. & Diamant, H. Law of corresponding states for osmotic swelling of vesicles. Soft Matter 8, 2185–2193 (2012).

47. Pilizota, T. & Shaevitz, J. W. Fast, multiphase volume adaptation to hyperosmotic shock by *Escherichia coli*. PLoS ONE 7, e35205 (2012).

48. Rojas, E., Theriot, J. A. & Huang, K. C. Response of *Escherichia coli* growth rate to osmotic shock. Proc. Natl. Acad. Sci. USA 111, 7807–7812 (2014).

49. Bendezú, F. O. & de Boer, P. A. J. Conditional lethality, division defects, membrane involution, and endocytosis in *mre* and *mrd* shape mutants of *Escherichia coli*. J. Bacteriol. 190, 1792–1811 (2008).

50. Reuter, M. et al. Mechanosensitive channels and bacterial cell wall integrity: does life end with a bang or a whimper? J. R. Soc. Interface 11, 20130850 (2013).

51. Bialecka-Fornal, M., Lee, H. J. & Phillips, R. The rate of osmotic downshock determines the survival probability of bacterial mechanosensitive channel mutants. J. Bacteriol. 197, 231–237 (2015).

52. Boer, M., Anishkin, A. & Sukharev, S. Adaptive MscS gating in the osmotic permeability response in *E. coli*: the question of time. Biochem. 50, 4087–4096 (2011).

53. Li, F., Chan, C. U. & Ohl, C. D. Yield strength of human erythrocyte membranes to impulsive stretching. Biophys. J. 105, 872–879 (2013).

54. Evans, E. A., Waugh, R. & Melnik, L. Elastic area compressibility modulus of red cell membrane. Biophys. J. 16, 585–595 (1976).

## Supplementary References

[1] Ventsel, E. & Krauthammer, T. Thin Plates and Shells: Theory, Analysis, and Applications (Marcel Dekker, Inc., 2001).

[2] Flügge, W. Stresses in Shells (Springer, 1960).

[3] Wong, F. et al. Mechanical strain-sensing implicated in cell shape recovery in *Escherichia coli*. Nat. Microbiol. 2, 17115 (2017).

[4] Bower, A. F. Applied Mechanics of Solids (CRC Press, 2010).

[5] Jiang, H. & Sun, S. X. Growth of curved and helical bacterial cells. Soft Matter 8, 7446–7451 (2012).

[6] Daly, K. E., Huang, K. C., Wingreen, N. S. & Mukhopadhyay, R. Mechanics of membrane bulging during cell-wall disruption in Gram-negative bacteria. Phys. Rev. E Stat. Nonlin. Soft Matter Phys. 83, 041922 (2011).

[7] Spurk, J. H. Fluid Mechanics: Problems and Solutions (Springer-Verlag, 1997).

